# Ghost in the Machine: Evidence for Non-Random Errors During Direct RNA Nanopore Sequencing Due to Post-Translocated RNA Folding

**DOI:** 10.64898/2025.12.02.691860

**Authors:** Jason M Needham, Philip Z Johnson, Anne E Simon

## Abstract

Direct RNA nanopore sequencing allows for the identification of full-length RNAs with a ∼10% error rate consisting of mismatches and small deletions. These errors are thought to be randomly distributed and structure-independent since RNA/cDNA duplexes are generated to prevent RNA structure formation prior to sequencing. When analyzing citrus yellow vein associated virus (CY1) reads during infection of *Nicotiana benthamiana,* viral (+/-)foldback RNAs (i.e., viral plus [+]-strands joined to [-]-strands) showed significantly higher error rates (mismatches and deletions) in the 5ʹ (+)RNA portion with errors that were relatively evenly distributed, while errors in the attached (-)RNA portion were less frequent and unevenly distributed. Non-foldback CY1 (+)RNAs from infected plants also showed an uneven distribution of errors, which correlated with errors in *in vitro* transcribed CY1 (+)RNA reads in both position and frequency. Hotspot errors in non-foldback CY1 (+)RNA and (-)RNA reads only weakly correlated, and hotspots were frequently located 5ʹ of known structural elements. Since nanopore sequencing is also used to identify RNA modifications, which depend on base-specific sequencing errors, algorithms for RNA modification detection were also examined for bias. We found that multiple programs predicted RNA modifications in *in vitro* transcribed CY1 RNA at the same positions and with similar confidence levels as with *in planta* CY1 RNA. These data suggest that direct RNA sequencing contains inherent error biases that may be associated with post-translocation RNA folding and low sequence complexity, and therefore extrapolations based on sequencing error require special consideration.

## INTRODUCTION

During nanopore sequencing, either DNA or RNA is directly sequenced by threading individual nucleic acids through a protein pore and measuring the change in voltage as each base translocates^1^. For direct RNA sequencing (DRS), the prior generation of an RNA/cDNA duplex is required to prevent secondary and tertiary RNA structures from forming prior to translocation that could vary the translocation speed^2^. Although more costly than traditional sequencing methods, DRS has many advantages. For example, amplification of the genetic material is not required^3,4^, and thus relative quantification of DNA and RNA is less affected by possible amplification bias^5^. Furthermore, the entire length of the nucleic acid is read continuously, allowing for confident assemblies of traditionally troublesome regions such as GC-rich and repetitive elements^6,7^. However, DRS results in a higher error rate than other sequencing methods^8^, which occurs when the voltage signature of a translocating base is not correctly identified. For example, an incorrect base may be called if the voltage too closely resembles another base, or if the base signal cannot be distinguished from the signal of the previous base. Alternatively, if a voltage shift occurs too rapidly to be identified, the base may be missed, resulting in an apparent deletion of the translocating base. While errors have been found to occur at higher frequency in homopolymeric stretches^9^, errors during nanopore sequencing are generally considered to be randomly distributed^10^.

Since each base is analyzed directly, DRS has also been reported to detect RNA base modifications^11,12^. Over 170 RNA modifications are known^13^, with functions that include crucial roles in RNA stability^14^, protein interaction^15^, translation efficiency^16^, and plant RNA intercellular movement^17,18^. While the gold standard for RNA modification detection is RNA pulldowns using antibodies specific to the modification^19^, this often deters exploratory analyses where RNA modification is not suspected. Antibody-based sequencing techniques can also fail to detect low-frequency modifications or produce nonspecific capture of unmodified RNA resulting in false positives. While antibody-independent assays have been developed using next-generation sequencing^19^, these techniques still include the inherent biases and limitations of short read sequencing. Thus, uncovering RNA modifications via DRS through detection of signal intensity perturbations caused by modified bases has the potential to significantly expand the identification and localization of RNA modifications in a wide variety of cellular and viral RNAs.

The advantages of DRS have been especially useful in the study of RNA viruses^20–23^. Umbra-like viruses (ULVs) are a newly discovered group of single-stranded plus-sense RNA ([+]RNA) viruses found in wild and agricultural plants worldwide^24^. While most plant viruses either encode movement proteins and one or more silencer suppressors, or are associated with a helper virus that provides those functions, some ULVs do not encode either a movement protein or silencer suppressors and are frequently found in plants without an accompanying virus^25^. The simplest ULV is citrus yellow-vein associated virus 1 (CY1; 2692 nt), which only encodes a replication-associated protein and the RNA-dependent RNA polymerase (RdRp) frameshift extension product, and yet is able to infect a variety of unrelated plants^26^. We recently leveraged the power of direct RNA nanopore sequencing to analyze the transcriptome of a CY1 infection in *Nicotiana benthamiana* and found evidence of undescribed long non-coding (lnc)RNAs, defective (D)-RNAs, and (+/-)foldback RNAs^2,27^. CY1 foldback RNAs are hypothesized to form during the transcription of plus-strand (+)RNA, when the RdRp releases the (-)RNA template but continues transcription using the newly synthesized (+)RNA as template, thus generating a single-stranded RNA with a 5ʹ (+)RNA segment attached to a nearly perfectly complementary 3ʹ (-)RNA segment^2,28^. RNA foldbacks, which are predicted to form long, dsRNA helical structures with a single-stranded apical loop, have unknown functions during viral infection.

For the current study, we further analyzed CY1 foldback RNAs, finding that the (+)RNA portions were basecalled with significantly higher mismatches and deletions than the already threaded (-)RNA portion in the same read. Additionally, non-foldback *in planta* (+)RNA reads and *in vitro* transcribed (IVT) (+)RNA reads showed a similar, nonrandom distribution of mismatches and deletions that were enriched in stretches of purines or pyrimidines and often appeared upstream of known stable RNA structures. Examination of IVT and *in planta* CY1 (+)RNA reads using a variety of RNA modification detection software predicted modifications at the exact same positions, and with similarly high confidences, despite IVT reads being devoid of any modifications. These data suggest that regions of low nucleotide complexity as well as post-translocation re-folding of RNA structures cause perturbations in the nanopore sequencing signal resulting in inherent, nonrandom basecalling errors. Furthermore, since RNA modification prediction software also relies on signal perturbation during DRS, these predictions should be viewed with caution without additional confirmatory evidence.

## MATERIALS AND METHODS

### Growth of N. benthamiana

Laboratory strain *N. benthamiana* seeds, originally collected by Benjamin Bynoe and housed at the Royal Botanic Gardens^29^, were initially seeded onto damp soil and germinated at 25°C with a 12 h light cycle and 70% humidity. After approximately 2 weeks, seedlings were transplanted into individual pots and grown at 25°C with a 12 h light cycle and 70% humidity until vacuum infiltration.

### Agroinfiltration of *N. benthamiana* with CY1

*Agrobacterium tumefaciens* strain GV3101 was transformed by electroporation with binary vector pCB301 containing full-length CY1 (NC_040311.1) immediately downstream of duplicated cauliflower mosaic virus 35S promoters and immediately upstream of a hammerhead ribozyme sequence. Transformed *A. tumefaciens* cultures were grown to an OD between 1.0 and 1.2 in 0.5 L of Luria-Bertani broth supplemented with antibiotics [rifampicin (20 µg/mL) and kanamycin (50 µg/mL)] over the course of ∼18 h, along with *A. tumefaciens* cultures transformed with standard RNA silencing suppressor p19. *A. tumefaciens* cultures were centrifuged at 5K rpm for 10 min using a Sorvall SLA-1500 rotor, resuspended in infiltration buffer (10 mM MgCl_2_; 10 mM MES; 100 ng/mL acetosyringone) at an OD of 1.2 for viral cultures and 0.4 for RNA silencing suppressor cultures, mixed in a 1:1 ratio of viral culture to RNA silencing suppressor culture, and incubated for 2 h at room temperature. *N. benthamiana* containing six true leaves were then submerged inverted in the mixed *A. tumefaciens* cultures and vacuum infiltrated using a negative pressure of -25 inHg for 30 sec. Plants were grown at 25°C with a 12 h light cycle. Systemic leaf and primary root stalk samples were harvested from infiltrated plants at 2 or 6 wpi.

### Extraction of RNA from infected plant samples

Total RNA was extracted from infected plant samples using 1 mL of TRIzol reagent (#15596026, Invitrogen) following the manufacturer’s instructions. Root samples were thoroughly ground with a mortar and pestle after being frozen in liquid nitrogen immediately prior to TRIzol extraction. Following TRIzol extraction, extracted root RNA samples were precipitated twice with equal volumes of 5M LiCl to remove excess polysaccharides in the RNA samples. All extracted RNA samples were purified using 65 μL of RNAClean XP beads (#A63987 Beckman Coulter) and analyzed by ethidium bromide-stained agarose gel electrophoresis prior to any downstream procedures.

### In vitro transcription of CY1 RNA

pET17b plasmid containing full-length CY1 gRNA sequence (GenBank: JX101610) immediately downstream of a T7 promoter was linearized with HindIII (#R0104M New England Biolabs) and used as template for in vitro transcription using T7 polymerase. In vitro transcribed CY1 gRNA sample volume was raised to 100 μL with ddH_2_O followed by addition of 100 μL of 5M LiCl and incubation at -20°C for 30 min. Samples were centrifuged at top speed for 30 min at 4°C followed by a 75% ethanol wash, air drying, and resuspension in ddH2O.

### Poly(A) tailing of RNA

Approximately 500 ng of RNA was mixed with ddH_2_O to a volume of 15.5 μL. Two microliters of 10X buffer (#B0276S New England Biolabs), 2 μL of 10 mM ATP (#B0756A New England Biolabs), and 0.5 μL of *E. coli* poly(A) polymerase (# M0276S New England Biolabs) were then added. Reactions were incubated at 37°C for 3 to 5 min and then terminated by addition of 5 μL of 50 mM EDTA. Poly(A) tailed RNA was purified using 65 μL of RNAClean XP beads (#A63987 Beckman Coulter) following manufacturer’s instructions and resuspended in 12 to 16 μL of ddH_2_O.

### Direct RNA and cDNA nanopore sequencing

For all DRS sequencing runs, sequencing libraries were prepared from poly(A)-tailed RNA samples using the direct RNA sequencing kit (SQK-RNA002) following manufacturer’s instructions and including the reverse transcription step to generate RNA/cDNA hybrids. Sequencing runs (6 to 18 h) were performed using version R9.4.1 flow cells and a MinION Mk1B device. Used flow cells were cleaned between runs using the flow cell wash kit (EXP-WSH004) following the manufacturer’s instructions. The MinKNOW desktop application (Oxford Nanopore) was used for basecalling of nanopore sequencing reads using the standard quality score threshold of 7 for direct RNA sequencing (corresponding to at least 80% read accuracy). For cDNA sequencing, 500 ng of poly-A tailed IVT CY1 RNA were used to generate a cDNA using oligo dT (IDT) using SuperScript III reverse transcriptase (#56575 Invitrogen) following the manufacturer’s instructions. The sequencing library was generated from the cDNA using the ligation sequencing kit (SQK-LSK114) and sequenced using a FLO-MIN114 R10 flowcell and a MinION Mk1B device. DNA reads were basecalled using the MinKNOW desktop application using the standard quality score threshold of 8 for DNA sequencing.

### Analysis of nanopore reads

Direct RNA and cDNA sequencing reads were aligned to the CY1 reference genome (NC_040311.1) or the *N. benthamiana* 5S ribosomal RNA (KP824744.1) using a locally run blast search (BLAST 2.12.0+) using the default parameters to a JSON output format expect for the foldback RNA, which were analyzed using the ‘blastn’ task parameter. JSON output files were analyzed using the custom analysis scripts deposited in the following GitHub repository: github.com/gr3nd31/Simon_lab/tree/main/nanopore_data_analysis. Using the blast alignments, positional abundance and relative error rates were calculated. Mismatch or deletion hotspots are defined as bases with error more than 2 standard deviations from the median IVT error, except for (-)RNA hotspots that are more than 2 standard deviations from the (-)RNA median. Data was visualized using R (4.4.3) in RStudio (2024.12.1.563). Error-prone positions were mapped onto structures published for CY1^26^ or 5S rRNA (http://combio.pl/rrna/).

### RNA modification analysis

Tentative RNA modification predictions were performed using tombo 1.5.1 (https://nanoporetech.github.io/tombo/), NanoPsu (https://github.com/sihaohuanguc/Nanopore_psU), and m6Anet v-2.1.0 (https://github.com/GoekeLab/m6anet) to detect 5mC, Ψ, and m6A, respectively. For each prediction algorithm, the developer’s protocol was followed.

### Statistical analysis and data availability

Statistical analyses were performed using R (4.4.3) in RStudio (2024.12.1.563) as described in the figure legends.

## RESULTS AND DISCUSSION

### CY1 foldback RNAs have an uneven distribution of errors

We previously reported that over 30% of the (-)-strand reads generated from DRS of samples from CY1-infected *N. benthamiana* were from (+/-)foldbacks, with the (-)RNA portion always downstream of the complementary (+)RNA^2^. While these (+/-)foldbacks varied in length (likely due to premature transcription termination by the RdRp during (+)RNA synthesis), the (-)RNA aligned portion consistently spanned 50% of the read (Fig. S1 A-C). While the (-)RNA portion of the (+/-)foldbacks aligned with high identity to CY1 reference sequence by BLASTn, the (+)RNA portion often failed to align unless a less stringent alignment algorithm was used (Fig. 1A, Fig. S1). The alignment failure was caused by the (+)RNA portion having an average mismatch/deletion frequency for all residues that was ∼3-fold higher than for the downstream (-)RNA portion (Fig. 1B and C, Fig. S1D and E). DRS has been reported to produce an error rate of ∼10%^30^, and all other CY1 (+)RNA reads (full-length gRNA, D-RNAs, lncRNAs, etc.) conformed to this error rate, as did reads generated from IVT CY1 gRNA (Fig. 1B and C). Thus, the elevated error rate in the (+)RNA portion of the (+/-)foldback was not due to any intrinsic characteristic of nanopore sequencing of CY1 (+)RNA.

**Figure 1.**
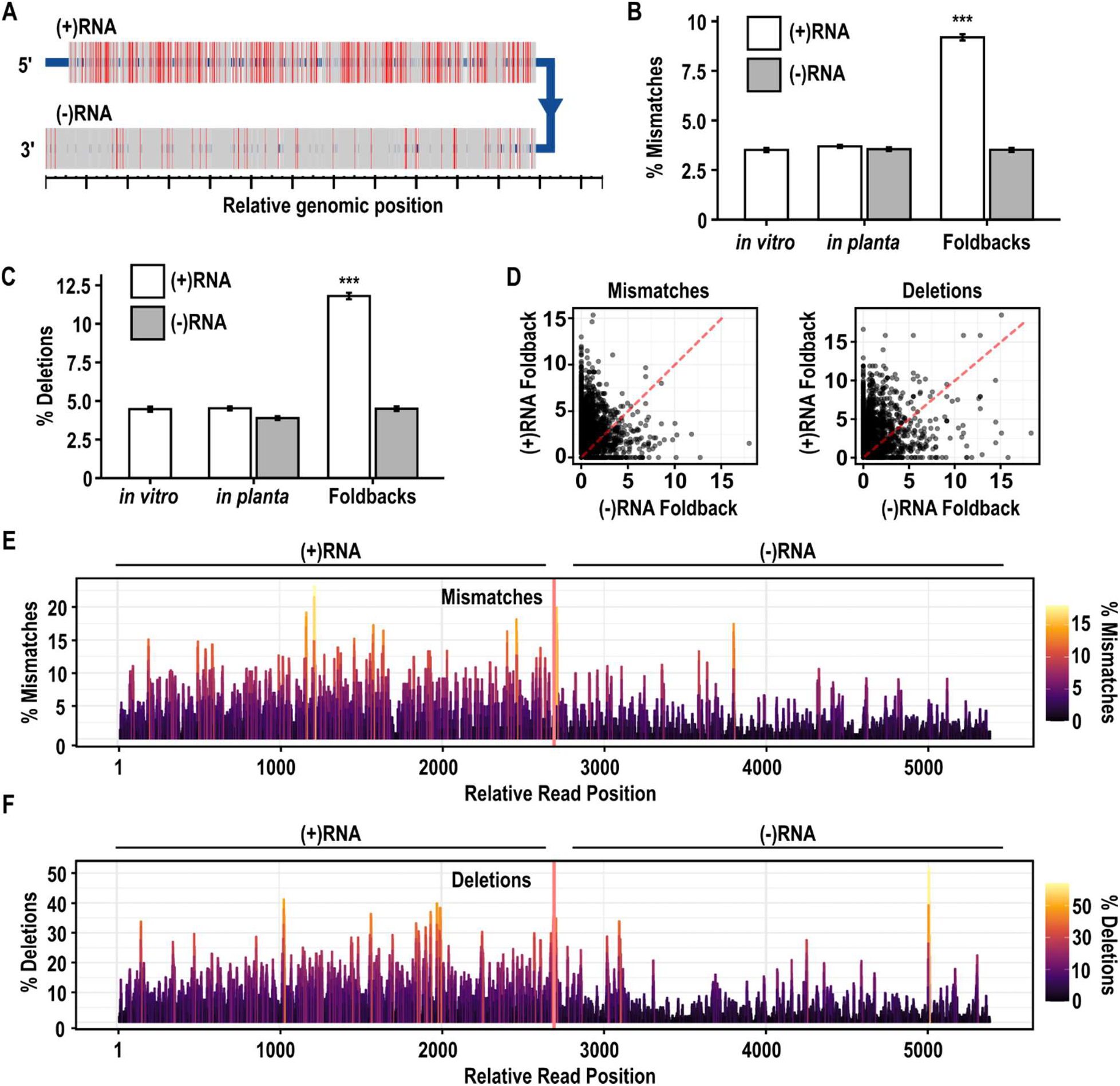
(+)RNA portions of CY1 foldback RNAs analyzed by DRS have a high error rate. (A) Representative (+/-)foldback CY1 sequences (n=87) collected from systemically infected leaves at 6-weeks post-infiltration (wpi). Nanopore reads that contained both (+)RNA and (-)RNA were aligned to the CY1 genome using BLASTn. Mismatches are in red and deletions are represented by gaps. (B) Average frequency of mismatches for residues in (+)RNA or (-)RNA reads from IVT CY1 RNA (*in vitro*), and (+/-)foldback (Foldback) and non-foldback reads from CY1-infected leaves (*in planta*). Error bars represent standard error and statistical significance from the *in vitro* sample was determined by one-way ANOVA and a Tukey post-hoc (*** : *p* < 0.001). (C) Average frequency of deletions in the reads described in (B). (D) Dot plot correlation of mismatch and deletion frequency at nucleotide positions in (+)RNA or (-)RNA aligned sequences in (+/-)foldback reads. Red dotted line represents a perfect 1:1 correlation. (E-F) Aggregate alignment of (+/-)foldback reads denoting mismatch frequency (E) or deletion frequency (F) normalized to the same position on the non-foldback reads in the same sample.

To determine if basecalling errors in the (+)RNA region of (+/-)foldbacks occurred at identical locations in the (-)RNA portions (i.e., the errors reflect natural *in vivo* generated alterations since the [+]RNA segment was the template for the [-]RNA segment), the error rate at each residue was normalized to non-foldback RNA. As shown in Fig. 1D, no correlation of mismatch/deletion errors was found between the (+)RNA and (-)RNA portions, indicating that errors were likely generated during sequencing. In addition, the (+)RNA portions showed a more even distribution of errors while distinct error hotspots were common for the (-)RNA portions (Fig. 1E and 1F). Whereas the average error rate for the (-)RNA portions was lower than the error rate for the (+)RNA portions, error hotspots within the (-)RNA portions occurred at similar rates as the average residue error rate for the (+)RNA portions. These findings suggest that a similar mechanism induced nanopore sequencing errors on both (+)RNA and (-)RNA portions of (+/-)foldbacks, but these errors occurred at a higher frequency consistently across the (+)RNA portion.

Single-stranded (+/-)foldback RNAs are unusual in that they fold into fully base-paired hairpins. Prior to nanopore sequencing, all intramolecular RNA structure is eliminated since RNAs are reverse transcribed to generate RNA/cDNA duplexes^10^. Once the RNA separates from the cDNA and is translocated through the membrane pore, intramolecular folding can then place. Since the (-)RNA portion of a (+/-)foldback is sequenced first, the translocated (-)RNA sequence should initially adopt structures comparable with non-foldback sequenced (-)RNA. However, once the complementary (+)RNA portion begins translocation, the foldback RNA should adopt a highly stable, double-stranded conformation. We hypothesize that the enhanced (+)RNA error rate is due to either torsional stress across the membrane pore or rapid shifts in translocation speed induced by the elongated dsRNA helix that forms when the (+)RNA segment translocates. Another possibility is that, similar to dsDNA, the dsRNA helix undergoes supercoiling^31^, which would induce novel torque and buckling pressures during (+)RNA basecalling. Any of these possibilities would account for both the higher mismatch/deletion frequencies during (+)RNA segment basecalling and the accompanying more uniform error distribution as the translocated RNA would be uniformly double-stranded.

### Non-foldback reads have a non-random distribution of mismatch/deletion errors

The apparently non-random nature of errors in the 3ʹ (-)RNA foldback segment suggested that initial secondary/tertiary structure folding within the translocated (-)RNA portions might affect the base-calling of nearby upstream nucleotides. If correct, then post-translocational RNA folding should also affect the sequencing error rate found for non-foldback RNAs, with hotspots occurring at specific locations upstream of local secondary/tertiary structure. Furthermore, these non-random hotspot errors should also be found at identical locations in the same RNA sequence generated by IVT.

To determine if non-random error hotspots occurred in non-foldback CY1 RNA reads, we analyzed CY1 reads from the same dataset as the (+/-)foldbacks as well as a dataset generated from DRS of IVT CY1 transcripts^2^. We selected reads that contained only a single (+)sense alignment and compared the mismatch and deletion error rates of individual bases between the two datasets (Fig. 2). A significant (r^2^ = 0.92, p < 0.001) positive correlation was found between the mismatch frequencies of *in planta* CY1 (+)RNA and IVT (+)CY1 RNA (Fig. 2A). While most bases were miscalled at a relatively low rate (median=1.8%, SD=3.5%), 130 bases of *in planta* CY1 (+)RNA possessed mismatch rates over 10%, which was more than 2 standard deviations from the average, and IVT (+)CY1 RNA sequences had similar mismatch rates for the same bases. We also compared the error rate of CY1 reads across multiple datasets generated from different tissues and from different timepoints post-infiltration and found remarkable consistencies in both the positions and rates of these errors (Fig. S2A and B).

**Figure 2.**
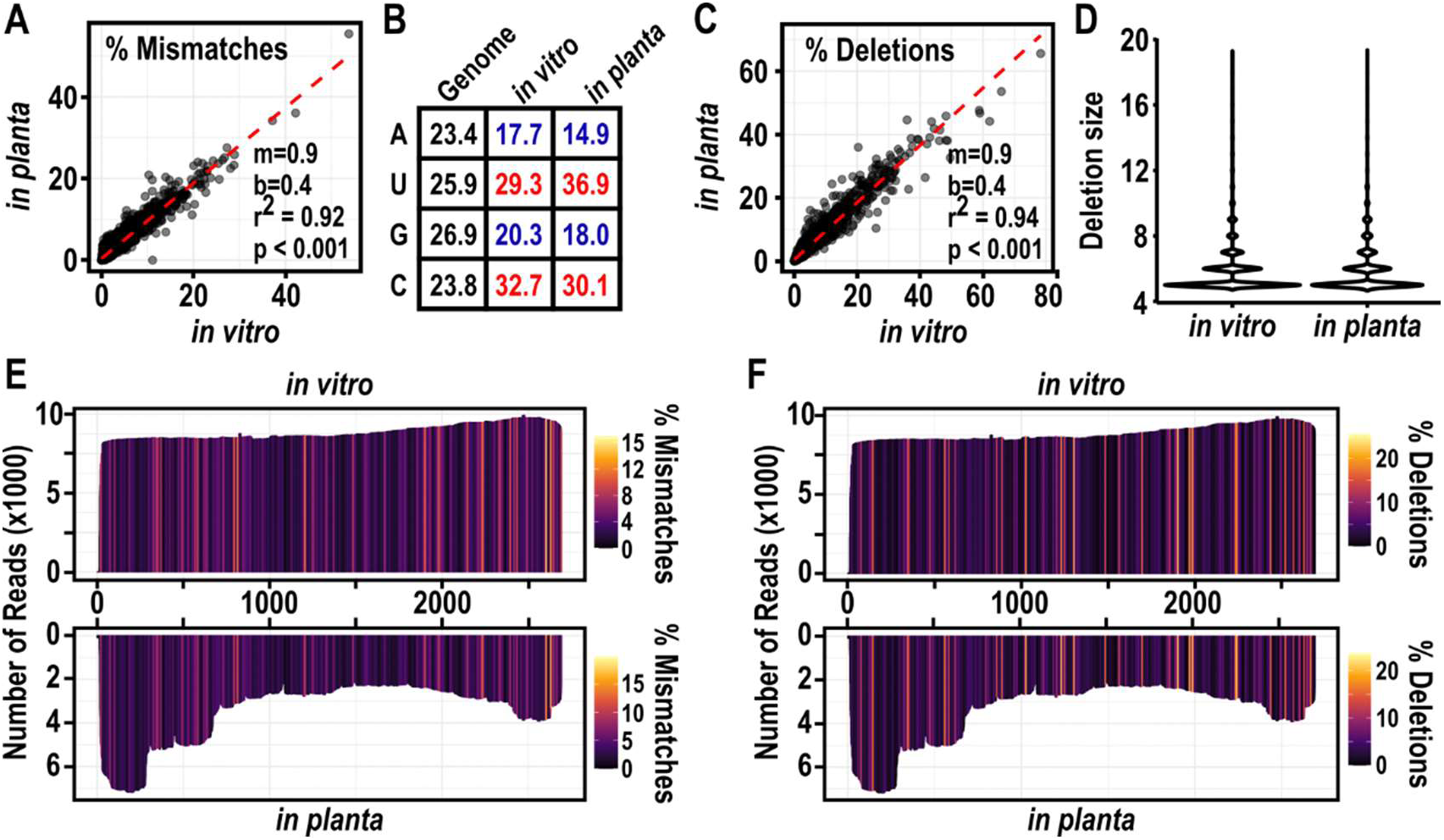
Mismatches and deletions reported for IVT and *in planta* CY1 RNA reads correlate in frequency and location. (A) Dot plot correlation of mismatch frequencies for each nucleotide of IVT RNA and RNA collected from 6 wpi leaves (*in planta*). Red dotted line represents a perfect 1:1 correlation and the statistics of the linear regression are shown. (B) The frequency of each nucleotide in CY1 gRNA (in black) compared to the frequency of that nucleotide being miscalled in *in vitro* or *in planta* (+)RNA reads (blue: less than expected based on composition, red: greater than expected based on composition). (C) Same correlation analysis as in (B) but comparing the deletion frequencies of each position within (+)RNA-aligned CY1 reads from *in vitro* and *in planta* reads. (D) Violin plot of the length of deleted stretches found for *in vitro* or *in planta* CY1-aligned reads. Since DRS frequently reports <4 nt deletions, deletions less than 4 nt in length were omitted from this analysis to enrich for less frequent deletion events. (E) Aggregate alignment of *in vitro* and *in planta* CY1-aligned reads colored by the average mismatch or deletion (F) frequency of a 4 nt sliding frame.

Both IVT and *in planta* reads showed a similar bias in the identity of miscalled bases, with cysteine (C) and uracil (U) more likely to be miscalled than adenine (A) and guanine (G), despite CY1 gRNA containing a relatively equal amount of each residue (Fig. 2B). Since a previous study found that A-to-G/G-to-A transversions were 3 to 5 times more likely in nanopore sequencing of DNA^9^, our finding that C-to-U/U-to-C transversion were more common in DRS suggests that the mismatch errors observed for DRS are distinct from known DNA nanopore sequencing biases.

Rates of deletions also strongly correlated between the IVT and *in planta* DRS reads, suggesting that deletion rates are also not randomly distributed across the CY1 sequence (Fig. 2C and F). As with mismatch errors, most bases exhibited a low deletion rate (median=1.9%, SD=6.5%), however the same182 positions in *in planta* and IVT (+)RNA possessed a deletion rate of greater than 15%. These same positions and deletion rates were present in *in planta* reads from multiple samples across time points and tissue types (Fig. S2C and D).

Previous studies reported that deletions in nanopore reads are frequently near homopolymer tracks, where inconsistent translocation speeds make it difficult to resolve the homopolymer^9^. Since homopolymer deletions are limited to the length of the homopolymer track and CY1 contains few homopolymer tracks longer than 3 nt, only deletions of greater than 4 nt were compared to reduce deletion hotspots that may be caused by homopolymer tracks and deletion hotspots occurring for other reasons (Fig. 2D). We found no differences in overall deletion size or in the distribution of deletion lengths between IVT and *in planta* CY1 RNA, suggesting that deletions in the reads resulted from nanopore error and were not legitimate deletions resulting from viral replication during infection. Altogether, these results strongly suggest that CY1 DRS mismatch and deletion sequencing errors are neither randomly distributed nor acquired during *in planta* infection and thus are intrinsic to the CY1 sequence.

### Error hotspots were frequently located upstream of known structural elements

High frequency error hotspots were identified as bases with local mismatch or deletion frequencies greater than 2 standard deviations from the median of all local frequencies (calculated from the IVT data). To better characterize the positions and sequence composition of these hotspots, the mismatch and deletion frequency for each residue in non-foldback CY1 RNAs was calculated by averaging a 5-nt sliding scale of mismatch/deletion frequencies (2 nt upstream and 2 nt downstream). As expected from the single nucleotide correlation plots (Fig. 2A and C), averaging local frequencies of mismatches (Fig. 2E) and deletions (Fig. 2F) showed a high correlation between IVT and *in planta* CY1 reads, with 22 hotspots for mismatches and 29 hotspots for deletions containing an average length of 7.8 ± 3 nt (Fig. 3). The average deletion rate for hotspots was higher than mismatch frequencies (17.9±3.9% vs 9.6±2.5%, respectively). While mismatch and deletion hotspots occasionally overlapped, different regions were more prone to generate either a mismatch or a deletion error during sequencing.

**Figure 3.**
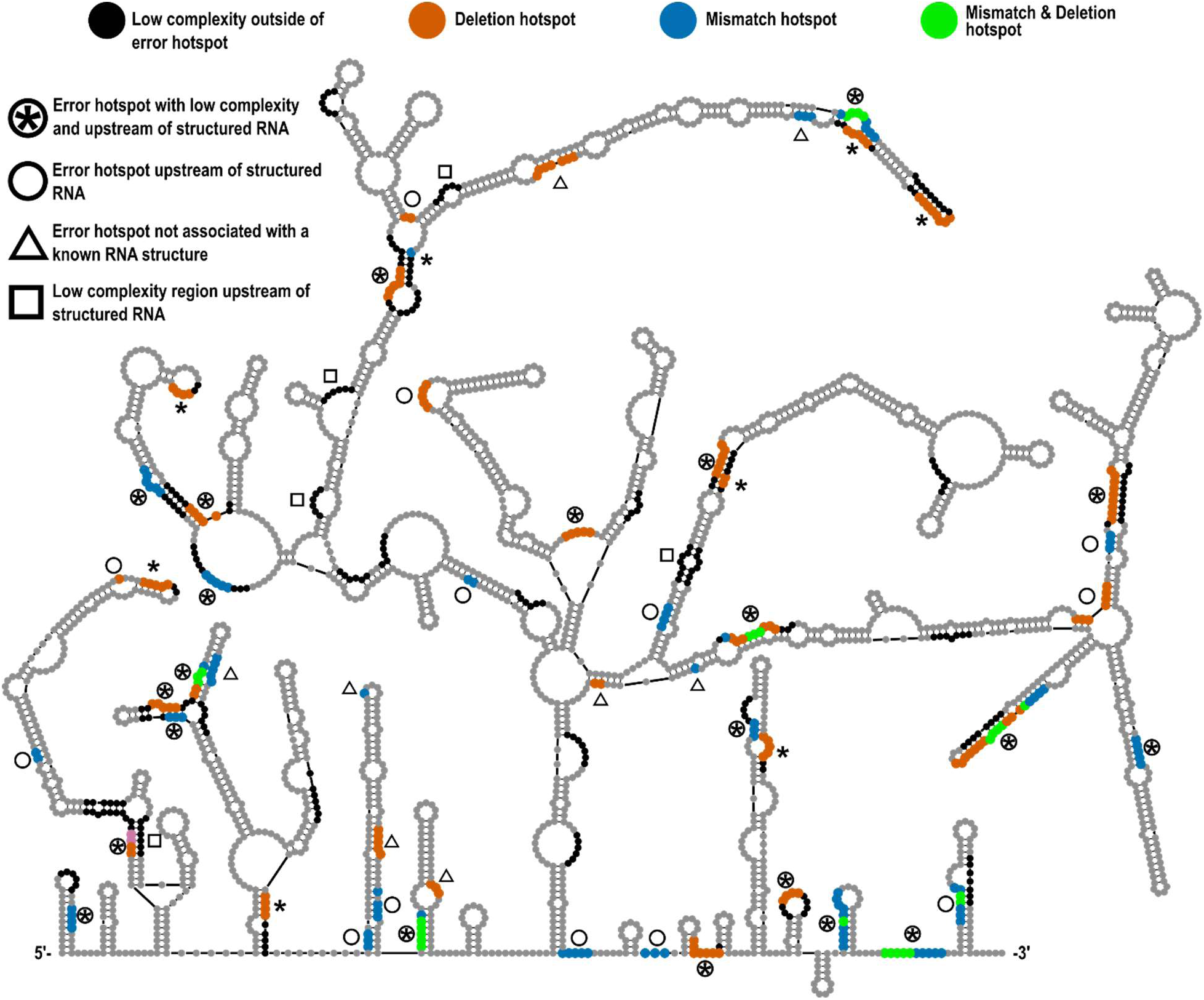
Deletion and mismatch hotspots mapped to CY1 secondary structure. Deletion and mismatch hotspots were identified by first averaging the deletion or mismatch frequency in a 4 nt sliding frame. Bases with an error rate greater than 2 standard deviations from the mean were classified as a deletion hotspot (orange), mismatch hotspot (blue) or both (green) and mapped back onto the solved secondary structure of CY1. Hotspots within a stretch of 6 or more purines/pyrimidine (low complexity regions) are indicated with an asterisk (*) for a pyrimidine stretch. Error hotspots that are less than 11 nucleotides upstream from a structure that could form during translocation are marked with a circle. In contrast, low complexity regions that were not associated with an error hotspots and were less than 11 nucleotides upstream from a structure are marked with a square. Error hotspots not associated with a known RNA structure or region of low complexity are marked with a triangle.

To determine if DRS hotspot errors in non-foldback reads were more common in purine or pyrimidine stretches, 4-nt stretches across the CY1 genome were evaluated, with 32% (867) of all possible windows consisting of consecutive purines or pyrimidines. Sixty three percent of these homopolymeric regions contained mismatch hotspots and 83% contained deletion hotspots, supporting the previous study’s findings^9^. However, many areas of low complexity did not present as error hotspots, suggesting that low complexity alone is insufficient and other parameters, including post-translocation structure folding, may also play a role. Since the elevated error rate of the (+/-)foldback RNA was only observed upstream of the proposed hairpin, we hypothesized that error hotspots arise from low complexity regions upstream of recently translocated RNA that folds into stable local structures. Such a mechanism is known to exist for bacterial terminators, where local hairpins form behind the RNA polymerase to terminate transcription at a downstream run of uracil residues^32^.

Error hotspots for non-foldback (+)RNA reads (31 mismatch and 32 deletion hotspots, a total of 51 unique hotspots) were mapped on the known secondary structure of CY1 (+)RNA^33,34^ (Fig. 3). Of the unique hotspots, 29 (57%) regions were within homopolymer stretches consisting of 6 or more purines or pyrimidines (Fig. 3, asterisk); 35 (69%) were within 11 nt upstream of a known structure that could be formed during translocation of the hotspot (Fig. 3, circles); and 22 (43%) were both within a homopolymer stretch and upstream of a known structure (Fig. 3, circles with asterisk). Although the CY1 genome contains 55 homopolymer stretches (Fig. 3, black bases or asterisk), only 5 (9%) of such stretches were upstream of a known structure and not associated with an error hotspot (Fig. 3, squares). In contrast, 15 (27%) of low complexity regions were neither upstream of a known structure nor associated with an error hotspot further supporting the model that low complexity sequence is insufficient to generate an error hotspot. Note that while RNA naturally folds co-transcriptionally 5ʹ to 3ʹ, post-translocation folding is 3ʹ to 5ʹ and thus some local structures will likely differ. Furthermore, structures may form transiently and not be present in the gRNA secondary structure map, which could explain the presence of error hotspots without a downstream structure (Fig. 3, triangles). Regardless, the association of the nonrandom mismatch and deletion hotspots with known structural elements in CY1, combined with data for the (+/-)foldbacks, suggest that DRS can be influenced by local folding of post-translocated 3ʹ RNA.

Aside from CY1 reads, the sequencing data also contained a high number of ribosomal RNAs, which are known to fold into stable structures necessary for their function^35,36^. As with CY1 reads, we found that 5S rRNA sequences had similar mismatch and deletion error rates between multiple datasets (Fig. S3). Many of these error hotspots also occurred near known structural elements, suggesting that 5S DRS errors are also nonrandomly distributed based in part on post-translocation folding. Like the CY1 error hotspots, mismatch and deletion hotspots did not occur at the same positions suggesting that these errors are generated by similar, yet distinct, mechanisms. A mismatch theoretically occurs when the electrical signal of the base is altered such that it mimics the signal of alternative bases. In contrast, a deletion likely occurs when the basecalling is unable to identify the signal transition between adjacent bases and thus only the initial base is identified. Therefore, it is likely that mismatch and deletion errors are associated with distinct sequence or structural elements that perturb base and transition signals, respectively. While the CY1 and 5S rRNA support this model, additional DRS data and solved structures are needed to further refine the hypothesis.

### cDNA sequencing does not recapitulate DRS error hotspots

Due to G/U and other non-canonical base-pairings, single-stranded (ss)RNA readily forms more complex secondary and tertiary structures than ssDNA^37^. If DRS error hotspots are due to folding of post-translocated RNA, then the distribution of errors from sequencing an RNA should differ from its cDNA version. Alternatively, if nanopore sequencing errors are solely a consequence of sequence complexity, then sequencing errors should correlate between (+)RNA, (-)RNA, and cDNA generated from IVT (+)RNA. To distinguish between these possibilities, cDNA was generated from IVT CY1 RNA using random hexamers and sequenced using the direct DNA ligation kit and a DNA flow cell. DRS reads of *in planta* CY1 from 6 weeks-post-infiltration (wpi) plants were split into (+)RNA and (-)RNA reads and compared with the positional mismatch/deletion error rates for the cDNA (Fig. 4). Unlike the high correlation of positional errors between *in planta* (+)RNA and IVT (+)RNA, there was no correlation between the mismatch error rates (Fig. 4A and 4B) nor deletion error rates (Fig. 4D and 4E) for *in planta* CY1 (+)RNA and (-)RNA, despite the RNA possessing similar regions of low complexity. Furthermore, there was no correlation between the error rate of the cDNA and the (-)RNA (Mismatches: Fig. 4A and 4C, Deletions: Fig. 4D and 4F). Since cDNA and (-)RNA share the same nucleotide complexity, this further supports the hypothesis that low nucleotide complexity alone is not causing error hotspots. While this distinction may be due to differences in DNA and RNA basecalling algorithms, the (+)RNA and (-)RNA also do not show the same error distributions despite possessing the same regions of low complexity and being generated from the same sequencing run (Fig. 4B and 4E). While cDNA and RNA sequences showed no correlation in positional error, a slight positive correlation was found between (-)RNA and (+)RNA positional errors (note linear regression marked by the hatched red lines in Fig. 4B and 4E relative to 4C and 4F, respectively), suggesting that some errors occur in similar regions. Interestingly, unique (+)RNA and (-)RNA error hotspots were found more than expected on opposing sides of complimentary regions (Fig. 4G and Fig. S4). Since DRS sequences RNA from the 3ʹ to 5ʹ direction, this further supports a model in which local folding of RNA post-translocation affects the sequencing fidelity of DRS.

**Figure 4.**
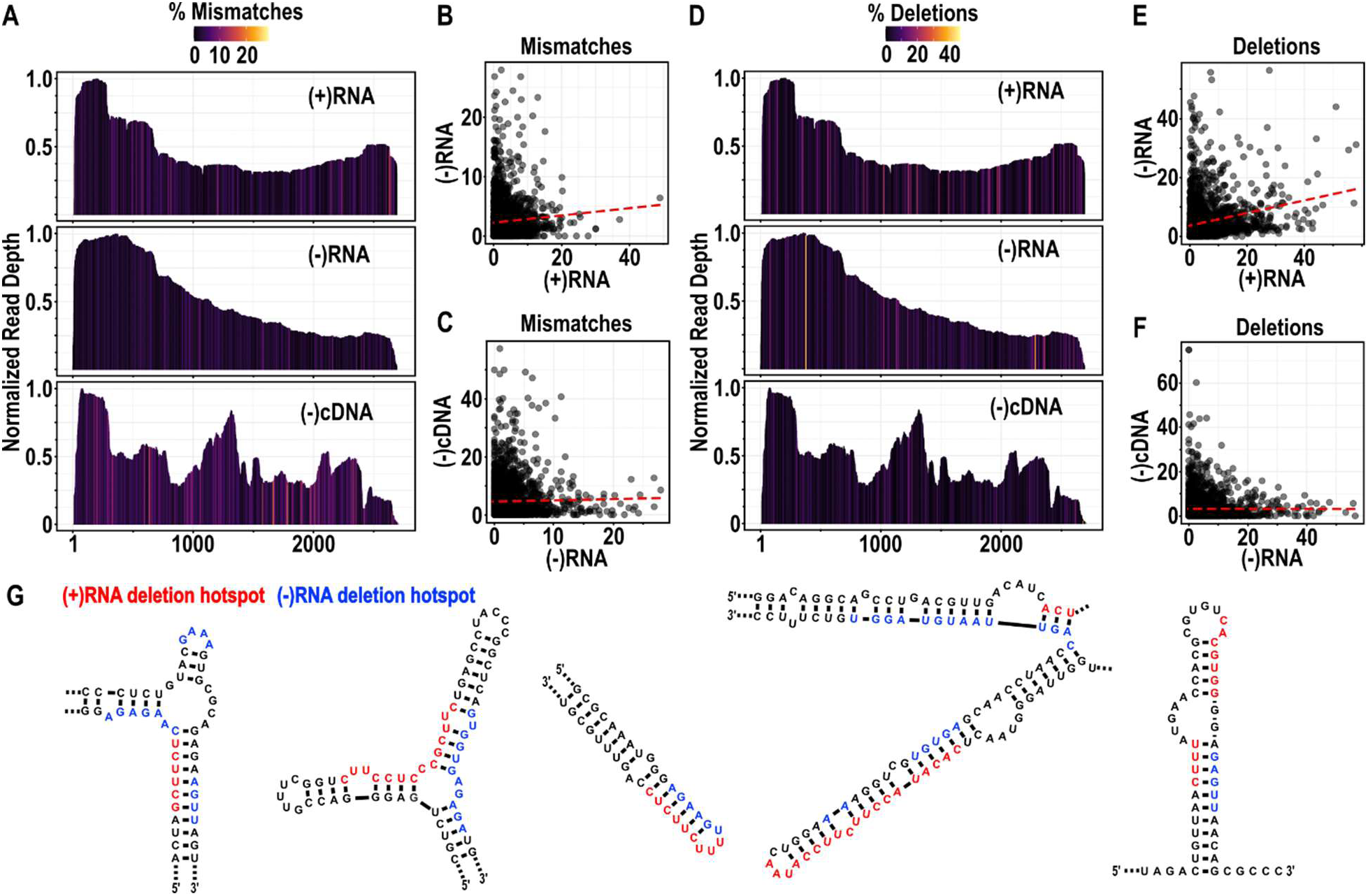
Error rates of (+)RNA and (-)RNA from DRS do not correlate with nanopore cDNA sequencing. (A) Aggregate alignment of the (+)RNA and (-)RNA reads from 6 wpi leaves and cDNA generated from reverse transcription of in-vitro transcribed CY1 (+)RNA using random hexamers. Positions are colored by the average mismatch frequency of a 4 nt sliding frame. (B) Dot plot and linear regression (red dotted line) of positions on the CY1 genome from (+)RNA and (-)RNA sequences shown in (A). (C) Dot plot and linear regression (red dotted line) of positions on the CY1 genome from (-)RNA and (-)cDNA sequences shown in (A). (D) Similar analysis as shown in (A) but colored by the average deletion frequency of a 4 nt sliding window. (E-F) Same dot plot and linear regression analysis as shown in (B-C) but comparing deletion frequencies. (G) Representative images of deletion hotspots for (+)RNA (red) and (-)RNA (blue) DRS reads mapped to secondary structure elements in the (+)CY1 genome.

### RNA modification detection programs did not distinguish between IVT and *in planta* CY1 (+)RNA reads

In addition to providing long read data, DRS is frequently used to predict DNA nucleotide modifications^38^ and, more recently, RNA modifications such as 5-methylcytosine (5mC), pseudouridine (Ψ), and N6-methyladenosine (m6A)^11,30^. These RNA modifications are predicted with high confidence in *in vivo* RNA samples by analyzing either the electrical signal generated during sequencing or the relative mismatch rates of specific nucleotides. Since mismatches are predicted based on variation in the electrical signal, and there is a high correlation between the mismatch frequencies of IVT and *in planta* (+)RNA CY1 reads (Fig. 2), we hypothesized that RNA modification detection programs may be incorrectly reporting modified nucleotides due to the nonrandom error distribution.

To determine if RNA modification software mis-identifies modified residues, we analyzed IVT and *in planta* (+)RNA reads for three modifications: 5mC using Tombo^39^ (Fig. 5A and B), Ψ using NanoPsu^40^ (Fig. 5C and D), and m6A using m6Anet^41^ (Fig. 5E and F). For each modification calling system, the IVT and 6 wpi CY1 reads were independently analyzed to predict modified residues.

**Figure 5.**
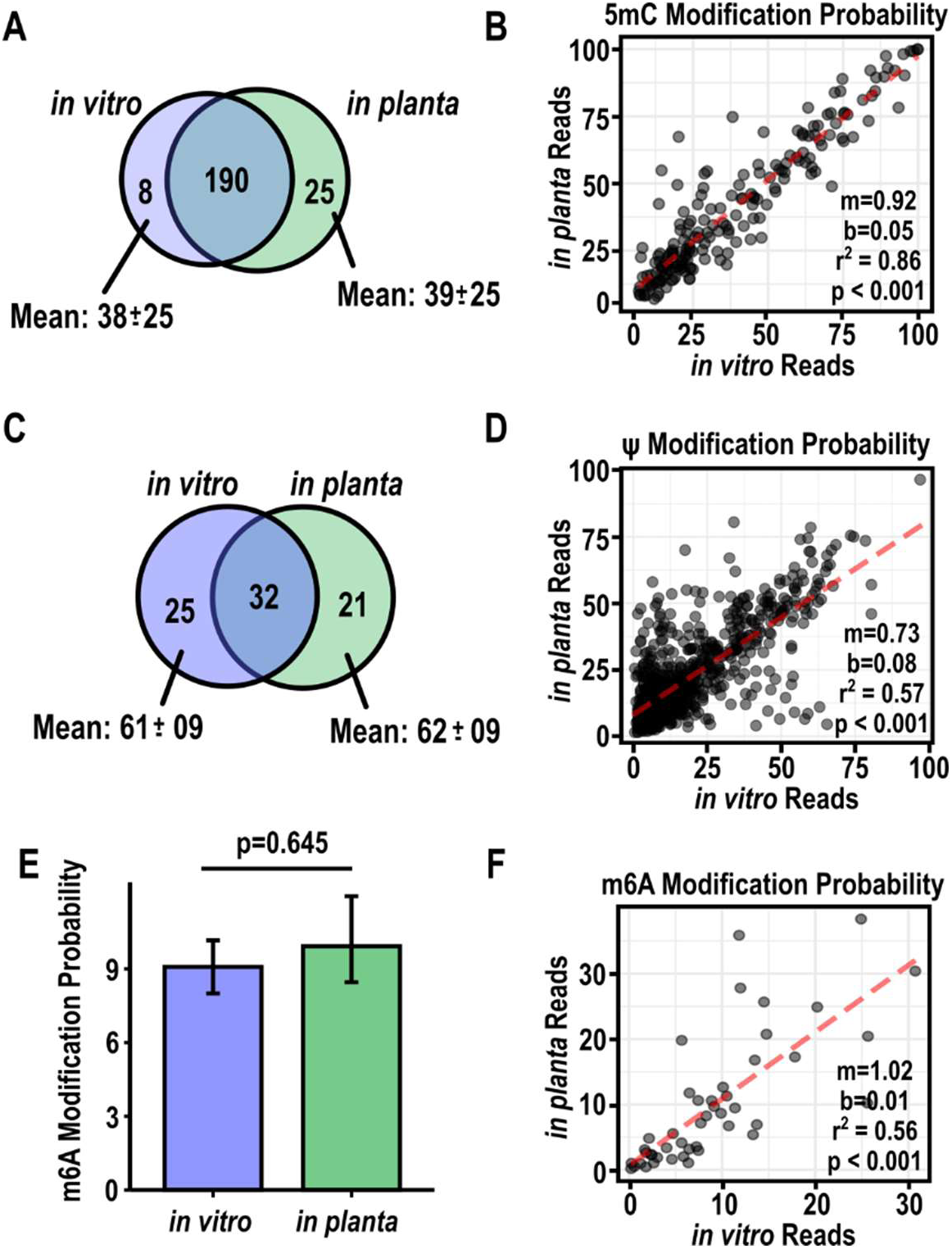
Predicted RNA modification in IVT CY1 reads mirror predictions in *in planta* reads. (A) Venn diagram of cytosine residues predicted to be methylated (5mC) in IVT (blue) or *in planta* (green) CY1 reads using Tombo. The mean 5mC probability and standard deviation for each set of cytosines is shown. (B) Dot plot and linear regression (red hatched line) of cytodine residues predicted to be 5mC. Statistics from a linear regression analysis are shown. (C) Venn diagram of predicted pseudouridine (Ψ) modifications in IVT (blue) or *in planta* (green) CY1 reads with probabilities at least 2 standard deviations above the mean modification probability. Mean Ψ probabilities and standard deviations for each set are indicated. (D) Dot plot and linear regression (red hatched line) of uracils predicted to be pseudouridines. Statistics from a linear regression analysis are shown. (E) Bar graph of the methylation probabilities of 46 adenosines in IVT and *in planta* reads in RRACH motifs. Error bars represent standard deviation and p-value was calculated by Student’s t-test. (F) Dot plot and linear regression (red hatched line) of methylation probabilities of adenosines within a RRACH motif in IVT or *in planta* CY1 reads. Statistics from a linear regression analysis are shown.

Within the IVT reads, which could not legitimately contain 5mC modifications, Tombo identified 198 residues as containing 5mC with an average probability of 38%, and 215 residues in the *in planta* reads with an average probability of 39% (Fig. 5A). Of these positions, a substantial majority (190) were identified in both the IVT and *in planta* reads. Direct comparison of the modification probability from the 190 positions identified in both sets of reads revealed a high correlation (Fig. 5B, r^2^ = 0.86, p <0.001). Although it is possible that the 25 positions identified only in the *in planta* reads may be legitimate 5mC modifications, the average probability of these *in planta* specific positions (19.2%) was both lower than the overall probability of the *in planta* positions and similar to the average probability of the IVT-specific positions (15.5%). While this suggests these *in planta*-specific positions are false-positives, we did note several positions identified in both the IVT and *in planta* reads that had a higher 5mC probability in the *in planta* reads than in the IVT reads. The high degree of correlation between IVT and *in planta* positions for predicted modifications, including several with confidence levels close to 100%, suggests that 5mC calling using Tombo is biased by nonrandom nanopore sequencing errors.

While Tombo directly analyzes electrical signals to predict 5mC modification, NanoPsu uses mismatch frequencies to calculate the probability of Ψ modification of uracils. Since NanoPsu generates modification probabilities for every uracil in an RNA, we limited our initial analysis to uracils with modification probabilities of at least 2 standard deviations from the mean (Fig. 5C). This resulted in a roughly equal number of hits in the IVT and *in planta* reads (57 and 53 hits, respectively) with 32 residues being selected in both read sets. Since the respective hit sets showed similar modification probabilities, we performed a linear regression analysis on all positions and found a strong positive correlation between the modification probabilities of IVT and *in planta* uracils (Fig. 5D, r^2^=0.57, p < 0.001) despite the lack of Ψ modification in IVT RNA. Although there were several positions in the *in planta* data with a higher probability than the same positions in the IVT data, the majority of Ψ probabilities demonstrated a strong correlation, suggesting that NanoPsu Ψ calling is also affected by nonrandom error.

As with Tombo, m6anet directly analyzes the electrical signal to predict m6A modifications but relies on Nanopolish to extract and analyze the electrical data. Since the RRACH motif for m6A modification has been well established, modification probabilities were only generated for the 46 possible RRACH motifs found in the CY1 genome^42^. Again, the overall probability of m6A modification was similar between the IVT and *in planta* reads (Fig. 5E), although the average probability of an m6A modification (∼9%) was much lower than that generated for 5mC (∼39%) and Ψ (∼20%). Linear regression analysis of m6A probabilities for IVF and *in planta* CY1 reads again showed a positive correlation (Fig. 5F, r^2^=0.56, p < 0.001). Altogether, these results suggest that all three modification detection software generate significant correlations for IVT and *in planta* CY1 RNA, indicating a strong bias in various modification-calling algorithms. Since we found that DRS results in consistent, nonrandom errors, we hypothesize that current base modification algorithms may fail to discriminate between legitimate RNA base modifications and nonrandom nanopore errors caused by properties intrinsic to the RNA.

### Conclusions

DRS is increasingly being used for long-read sequencing and identification of RNA base modifications^11,19^. Although DRS is associated with a 10% error rate, these errors are generally considered to be random with the exception of a general prevalence for homopolymeric stretches^9^. We show here that CY1 (+/-)foldbacks identified during DRS of infected *N. benthamiana* have an elevated frequency of mismatches and deletions in the 5ʹ (+)RNA segment only. This suggests that the increased error rate is driven by the fully complimentary structure of the foldback RNA. This hypothesis is supported by the observation that non-foldback CY1 RNAs from multiple sequencing runs have comparable error rates for the same nucleotides regardless of IVT or *in planta* generation. Furthermore, nucleotides with the highest error rates were frequently found in purine or pyrimidine stretches just upstream of known RNA structures. We therefore propose a model whereby transient torsional stress of local stable RNA structures that form just after translocation contribute to upstream sequencing errors by affecting either the electrical signal or translocation speed (Fig. 6A and B). While this stress may contribute generally to errors in upstream sequences, problematic sequences such as homopolymer stretches would be especially prone to increased error during basecalling. For a perfectly complementary sequence, such as the CY1 foldbacks, this results in an increased rate of evenly distributed errors during basecalling of the 5ʹ portion (Fig. 6C). While DRS technology possesses many benefits over traditional or next generation sequencing, these findings suggest caution in drawing conclusions from DRS data based solely on error rates as currently used by some RNA modification prediction programs.

**Figure 6.**
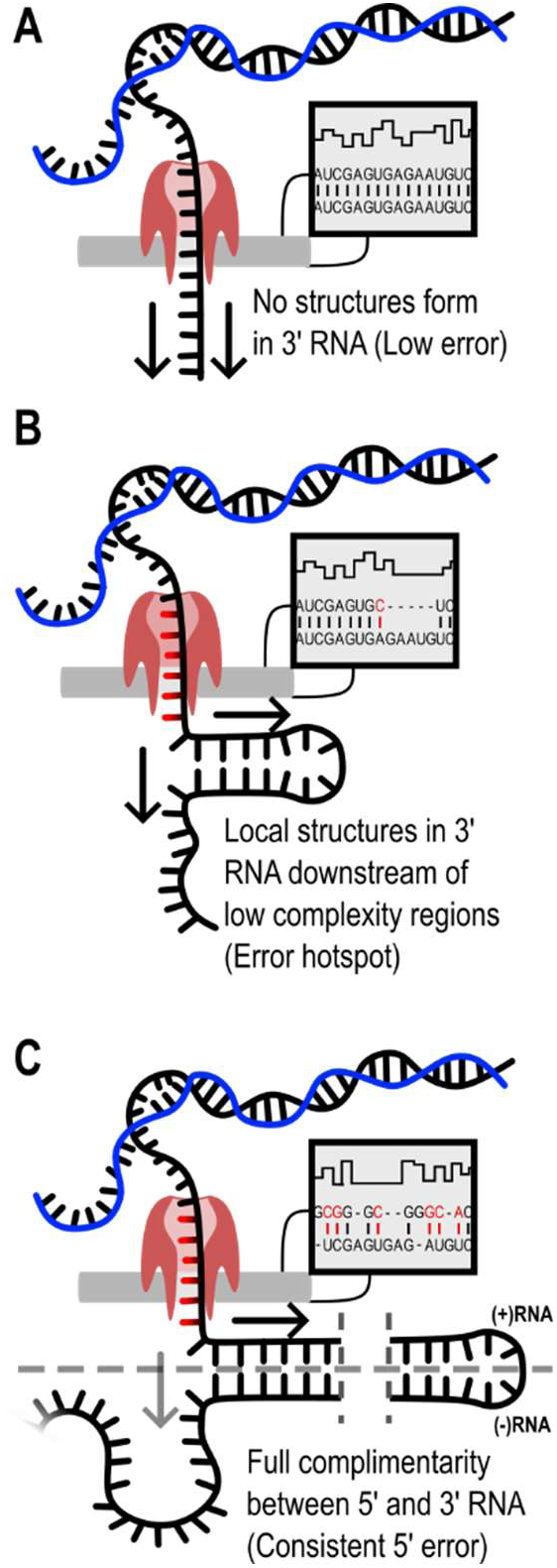
Model of 3ʹ structure-induced Nanopore error. (A) RNA/cDNA duplex (black and blue helix) prevents secondary structures in an RNA from interfering with translocation. Duplex is unwound before translocation and the RNA translocates through the pore in the 3ʹ to 5ʹ direction. Electrical signals are basecalled and aligned to the reference genome with a relatively low frequency and errors are evenly distributed across the read if translocated RNA does not quickly fold into a stable structure. (B) If the just translocated RNA forms a stable local structure, this results in torsional stress and/or alters translocation speed, which impairs basecalling and results in a nonrandom increase in mismatch and deletion errors upstream of these elements. We suggest that these errors are more likely if a stretch of purines or pyrimidines is just upstream of the structure. (C) During the sequencing of (+/-)foldback RNAs, basecalling of the initial (-)RNA sequence is only affected by local structure. After the complementary (+)RNA sequence begins to translocate through the pore, the (-)RNA base pairs with the (+)RNA to form dsRNA (transition point between the (-)RNA and the (+)RNA is marked by grey dashed line). This results in constant torsional stress and/or translocation speed, which is basecalled as an even distribution of elevated mismatch and deletion errors.

## ADDITIONAL INFORMATION

### Author Contributions

JM Needham contributed to this work through conceptualization, data curation, software development, formal analysis, validation, investigation, visualization, methodology, writing the original draft and editing. PZ Johnson contributed by investigation of the original dataset. AE Simon contributed by providing resources, supervision, funding acquisition, project management, and reviewing and editing of the manuscript.

### Data Availability

Datasets used to generate figures can be found at https://zenodo.org/records/15255459 and scripts used to generate figures from the nanopore sequencing data can be found at https://github.com/gr3nd31/Simon_lab/tree/main/nanopore_data_analysis.

## References

1. Rhee M, Burns MA. Nanopore sequencing technology: research trends and applications. Trends Biotechnol 2006; 24:580–6.

2. Johnson PZ, Needham JM, Lim NK, Simon AE. Direct nanopore RNA sequencing of umbra-like virus-infected plants reveals long non-coding RNAs, specific cleavage sites, D-RNAs, foldback RNAs, and temporal- and tissue-specific profiles. NAR Genom Bioinform 2024; 6:lqae104.

3. Sanger F, Coulson AR. A rapid method for determining sequences in DNA by primed synthesis with DNA polymerase. J Mol Biol 1975; 94:441–8.

4. Bustin SA. Absolute quantification of mRNA using real-time reverse transcription polymerase chain reaction assays. J Mol Endocrinol 2000; 25:169–93.

5. Martínez M, Harms L, Abele D, Held C. Mitochondrial Heteroplasmy and PCR Amplification Bias Lead to Wrong Species Delimitation with High Confidence in the South American and Antarctic Marine Bivalve Aequiyoldia eightsii Species Complex. Genes (Basel) 2023; 14:935.

6. Warburton PE, Sebra RP. Long-Read DNA Sequencing: Recent Advances and Remaining Challenges. Annu Rev Genomics Hum Genet 2023; 24:109–32.

7. Nomburg J, Zou W, Frost TC, Datta C, Vasudevan S, Starrett GJ, Imperiale MJ, Meyerson M, DeCaprio JA. Long-read sequencing reveals complex patterns of wraparound transcription in polyomaviruses. PLOS Pathogens 2022; 18:e1010401.

8. Liu-Wei W, van der Toorn W, Bohn P, Hölzer M, Smyth RP, von Kleist M. Sequencing accuracy and systematic errors of nanopore direct RNA sequencing. BMC Genomics 2024; 25:528.

9. Delahaye C, Nicolas J. Sequencing DNA with nanopores: Troubles and biases. PLOS ONE 2021; 16:e0257521.

10. Marinov GK. On the design and prospects of direct RNA sequencing. Briefings in Functional Genomics 2017; 16:326–35.

11. Abebe JS, Verstraten R, Depledge DP. Nanopore-Based Detection of Viral RNA Modifications. mBio 2022; 13:e03702–21.

12. Zhao L-Y, Song J, Liu Y, Song C-X, Yi C. Mapping the epigenetic modifications of DNA and RNA. Protein & Cell 2020; 11:792–808.

13. Boccaletto P, Stefaniak F, Ray A, Cappannini A, Mukherjee S, Purta E, Kurkowska M, Shirvanizadeh N, Destefanis E, Groza P, et al. MODOMICS: a database of RNA modification pathways. 2021 update. Nucleic Acids Res 2022; 50:D231–5.

14. Boo SH, Kim YK. The emerging role of RNA modifications in the regulation of mRNA stability. Exp Mol Med 2020; 52:400–8.

15. Zaccara S, Ries RJ, Jaffrey SR. Reading, writing and erasing mRNA methylation. Nat Rev Mol Cell Biol 2019; 20:608–24.

16. Shi H, Chai P, Jia R, Fan X. Novel insight into the regulatory roles of diverse RNA modifications: Re-defining the bridge between transcription and translation. Molecular Cancer 2020; 19:78.

17. Maizel A, Markmann K, Timmermans M, Wachter A. To move or not to move: roles and specificity of plant RNA mobility. Current Opinion in Plant Biology 2020; 57:52–60.

18. Wilkinson E, Cui Y-H, He Y-Y. Roles of RNA Modifications in Diverse Cellular Functions. Front Cell Dev Biol 2022; 10:828683.

19. Zhang Y, Lu L, Li X. Detection technologies for RNA modifications. Exp Mol Med 2022; 54:1601–16.

20. Xu L, Berninger A, Lakin SM, O’Donnell V, Pierce JL, Pauszek SJ, Barrette RW, Faburay B. Direct RNA Sequencing of Foot-and-mouth Disease Virus Genome Using a Flongle on MinION. Bio Protoc 2024; 14:e5017.

21. Donovan-Banfield I, Milligan R, Hall S, Gao T, Murphy E, Li J, Shawli GT, Hiscox J, Zhuang X, McKeating JA, et al. Direct RNA sequencing of respiratory syncytial virus infected human cells generates a detailed overview of RSV polycistronic mRNA and transcript abundance. PLoS One 2022; 17:e0276697.

22. Viehweger A, Krautwurst S, Lamkiewicz K, Madhugiri R, Ziebuhr J, Hölzer M, Marz M. Direct RNA nanopore sequencing of full-length coronavirus genomes provides novel insights into structural variants and enables modification analysis. Genome Res 2019; 29:1545–54.

23. Keller MW, Rambo-Martin BL, Wilson MM, Ridenour CA, Shepard SS, Stark TJ, Neuhaus EB, Dugan VG, Wentworth DE, Barnes JR. Direct RNA Sequencing of the Coding Complete Influenza A Virus Genome. Sci Rep 2018; 8:14408.

24. Simon AE, Quito-Avila DF, Bera S. Expanding the Plant Virome: Umbra-Like Viruses Use Host Proteins for Movement. 2024 [cited 2024 Jul 31]; Available from: https://www.annualreviews.org/content/journals/10.1146/annurev-virology-111821-122718

25. Ying X, Bera S, Liu J, Toscano-Morales R, Jang C, Yang S, Ho J, Simon AE. Umbravirus-like RNA viruses are capable of independent systemic plant infection in the absence of encoded movement proteins. PLOS Biology 2024; 22:e3002600.

26. Kwon S-J, Bodaghi S, Dang T, Gadhave KR, Ho T, Osman F, Al Rwahnih M, Tzanetakis IE, Simon AE, Vidalakis G. Complete Nucleotide Sequence, Genome Organization, and Comparative Genomic Analyses of Citrus Yellow-Vein Associated Virus (CYVaV). Front Microbiol 2021; 12:683130.

27. Johnson PZ. Translation, Replication and Transcriptomics of the Simplest Plus-Strand RNA Plant Viruses [Internet]. ProQuest Dissertations and Theses2024; Available from: https://www.proquest.com/dissertations-theses/translation-replication-transcriptomics-simplest/docview/3108156930/se-2?accountid=14696

28. Richards OC, Hey TD, Ehrenfeld E. Poliovirus snapback double-stranded RNA isolated from infected HeLa cells is deficient in poly(A). J Virol 1987; 61:2307–10.

29. Goodin MM, Zaitlin D, Naidu RA, Lommel SA. Nicotiana benthamiana: Its History and Future as a Model for Plant–Pathogen Interactions. MPMI 2008; 21:1015–26.

30. Jain M, Abu-Shumays R, Olsen HE, Akeson M. Advances in nanopore direct RNA sequencing. Nat Methods 2022; 19:1160–4.

31. Lipfert J, Skinner GM, Keegstra JM, Hensgens T, Jager T, Dulin D, Köber M, Yu Z, Donkers SP, Chou F-C, et al. Double-stranded RNA under force and torque: Similarities to and striking differences from double-stranded DNA. Proceedings of the National Academy of Sciences 2014; 111:15408–13.

32. Santangelo TJ, Artsimovitch I. Termination and antitermination: RNA polymerase runs a stop sign. Nat Rev Microbiol 2011; 9:319–29.

33. Liu J, Carino E, Bera S, Gao F, May JP, Simon AE. Structural Analysis and Whole Genome Mapping of a New Type of Plant Virus Subviral RNA: Umbravirus-Like Associated RNAs. Viruses 2021; 13:646.

34. Johnson PZ, Simon AE. RNAcanvas: interactive drawing and exploration of nucleic acid structures. Nucleic Acids Research 2023; 51:W501–8.

35. Vanden Broeck A, Klinge S. Eukaryotic Ribosome Assembly. Annu Rev Biochem 2024; 93:189–210.

36. Jang C, Needham JM, Johnson PZ, Gao F, Simon AE. Hairpin inserts in viral genomes are stable when they conform to the thermodynamic properties of viral RNA substructures. Journal of Virology 2025; 0:e01919–24.

37. Halder S, Bhattacharyya D. RNA structure and dynamics: A base pairing perspective. Progress in Biophysics and Molecular Biology 2013; 113:264–83.

38. Lau BT, Almeda A, Schauer M, McNamara M, Bai X, Meng Q, Partha M, Grimes SM, Lee H, Heestand GM, et al. Single-molecule methylation profiles of cell-free DNA in cancer with nanopore sequencing. Genome Medicine 2023; 15:33.

39. Stoiber M, Quick J, Egan R, Eun Lee J, Celniker S, Neely RK, Loman N, Pennacchio LA, Brown J. *De novo* Identification of DNA Modifications Enabled by Genome-Guided Nanopore Signal Processing. bioRxiv 2017; :094672.

40. Huang S, Zhang W, Katanski CD, Dersh D, Dai Q, Lolans K, Yewdell J, Eren AM, Pan T. Interferon inducible pseudouridine modification in human mRNA by quantitative nanopore profiling. Genome Biology 2021; 22:330.

41. Hendra C, Pratanwanich PN, Wan YK, Goh WSS, Thiery A, Göke J. Detection of m6A from direct RNA sequencing using a multiple instance learning framework. Nat Methods 2022; 19:1590–8.

42. Wang K, Peng J, Yi C. The m6A Consensus Motif Provides a Paradigm of Epitranscriptomic Studies. Biochemistry 2021; 60:3410–2.

