## Supplemental Figures for "Ghost in the Machine: Evidence for Non-Random Errors During Direct RNA Nanopore Sequencing Due to Post-Translocated RNA Folding" (from another version of this preprint: version 1, 2025-12-04)

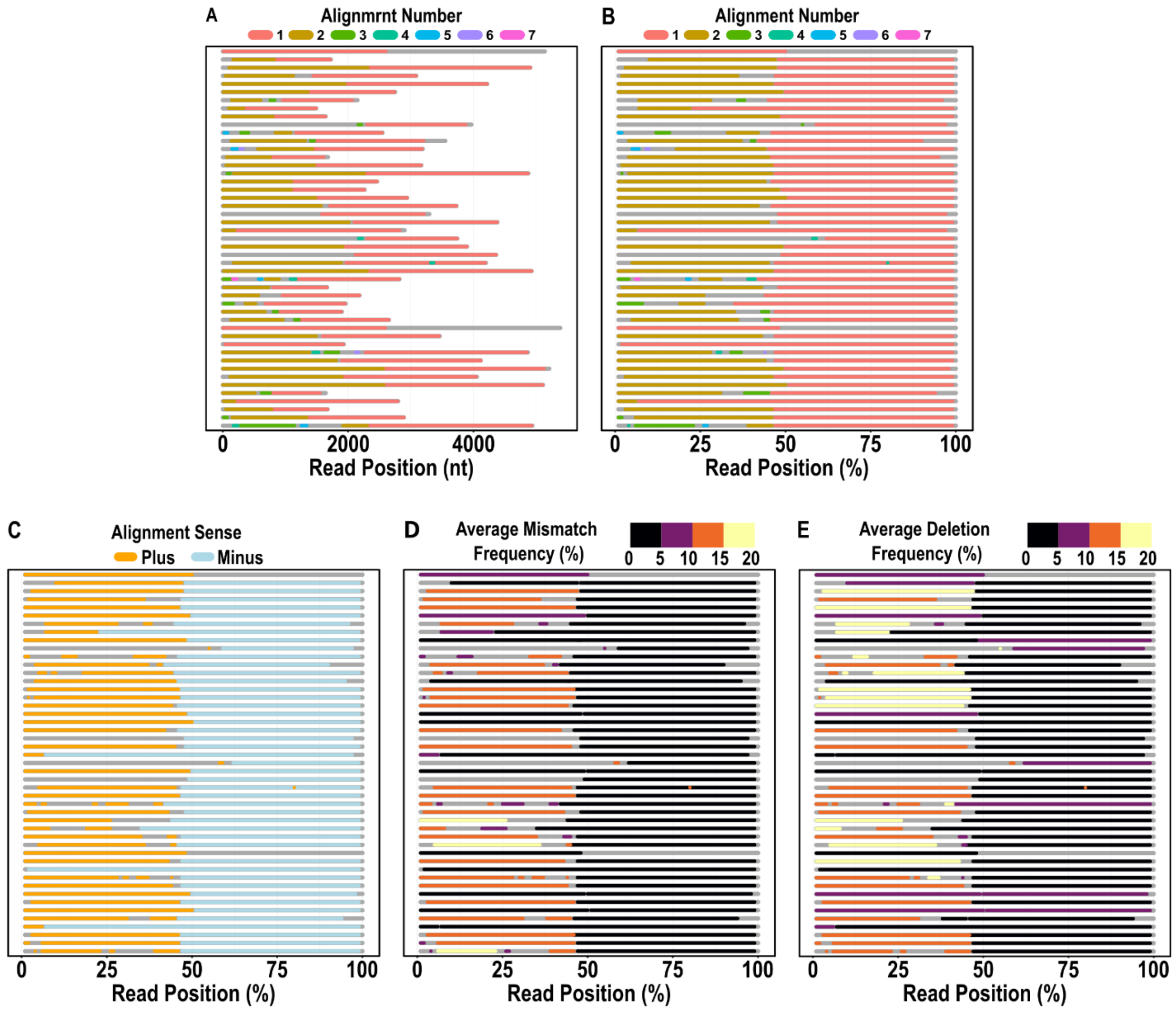


**Supplemental Figure 1. CY1 foldback RNA shows higher show higher error rates on the 5', (+)RNA region. (A)** Graphical representation of 47 foldback reads from 6 wpi CY1-infected *N. benthamiana* that aligned to at least 1500 bases in the CY1 genome. Read lengths are shown on the x-axis and are represented by gray bars with distinct alignment regions colored. Reads were aligned using BLASTn due to the high error rate in the 5' (+)RNA regions. **(B)** The same reads and alignment coloring shown in **(A)** but with read length normalized to 100%. **(C-E)** The same reads normalized to read length as in **(B)** but colored by **(C)** alignment, **(D)** average mismatch rate in the alignment region, or **(E)** the average deletion rate of the aligned region.

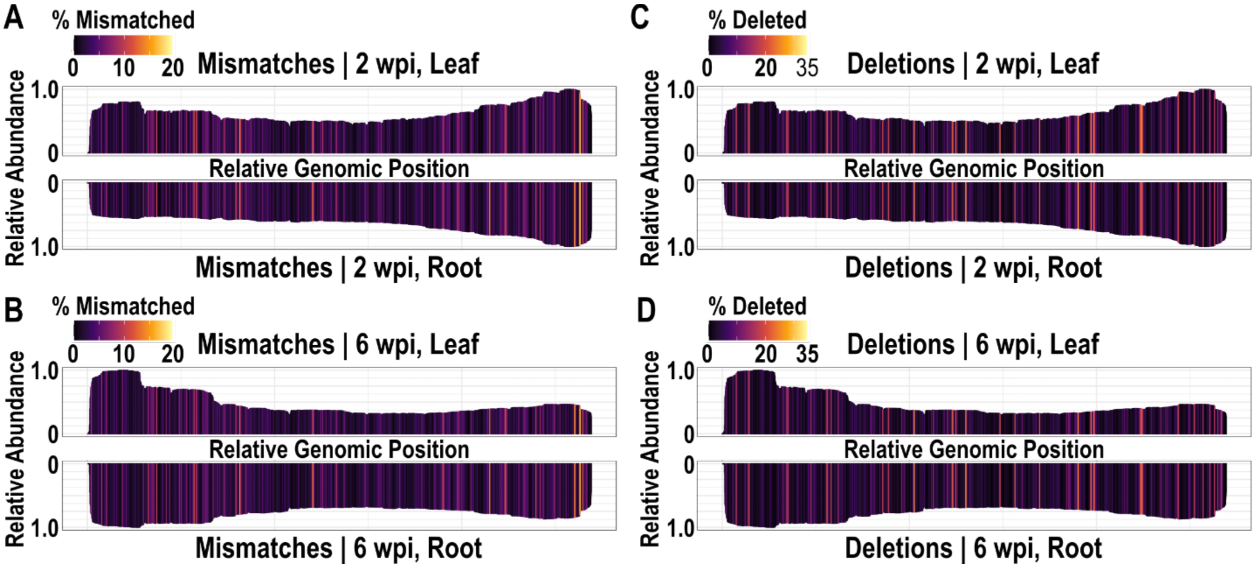


**Supplemental Figure 2. CY1 error hotspots correlate across multiple samples regardless of tissue or timepoint. (A-B)** Aggregate alignment of *in planta* CY1-aligned read from the systemic leaf (top) or root (bottom) collected at **(A)** 2 wpi and **(B)** 6 wpi. Positions are colored by the mismatch frequency calculated using a 4 nt sliding frame. **(C-D)** Same analysis as **(A-B)** but with positions colored by deletion frequency as calculated using a 4 nt sliding frame.

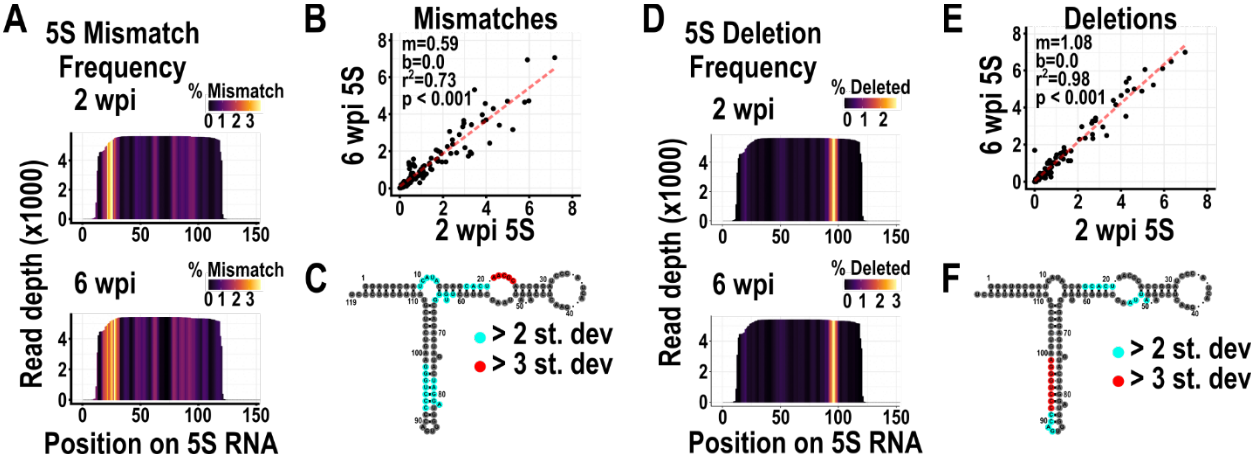


**Supplemental Figure 3. Non-viral, structured RNA show consistent, non-random errors.** **(A)** Aggregate alignment of systemic leaf reads collected at 2 and 6 wpi and aligned to the *N. benthamiana* 5S rRNA sequence and colored by mismatch frequency calculated using a 4 nt sliding frame. **(B)** Dot plot correlation of the mismatch frequencies found on 5S aligned reads from 2 wpi and 6 wpi samples. Red, dotted line represents a 1:1 correlation and statistics of the linear regression are shown. **(C)** The secondary structure of 5S rRNA with positions colored based on the mismatch frequencies. Positions with a mismatch frequency greater than 2 standard deviations from the mean 5S mismatch rate are in teal and positions with a mismatch frequency greater than 3 standard deviations are in red. **(D-F)** The same analysis as performed in **(A-C)** but colored based on deletion frequencies.

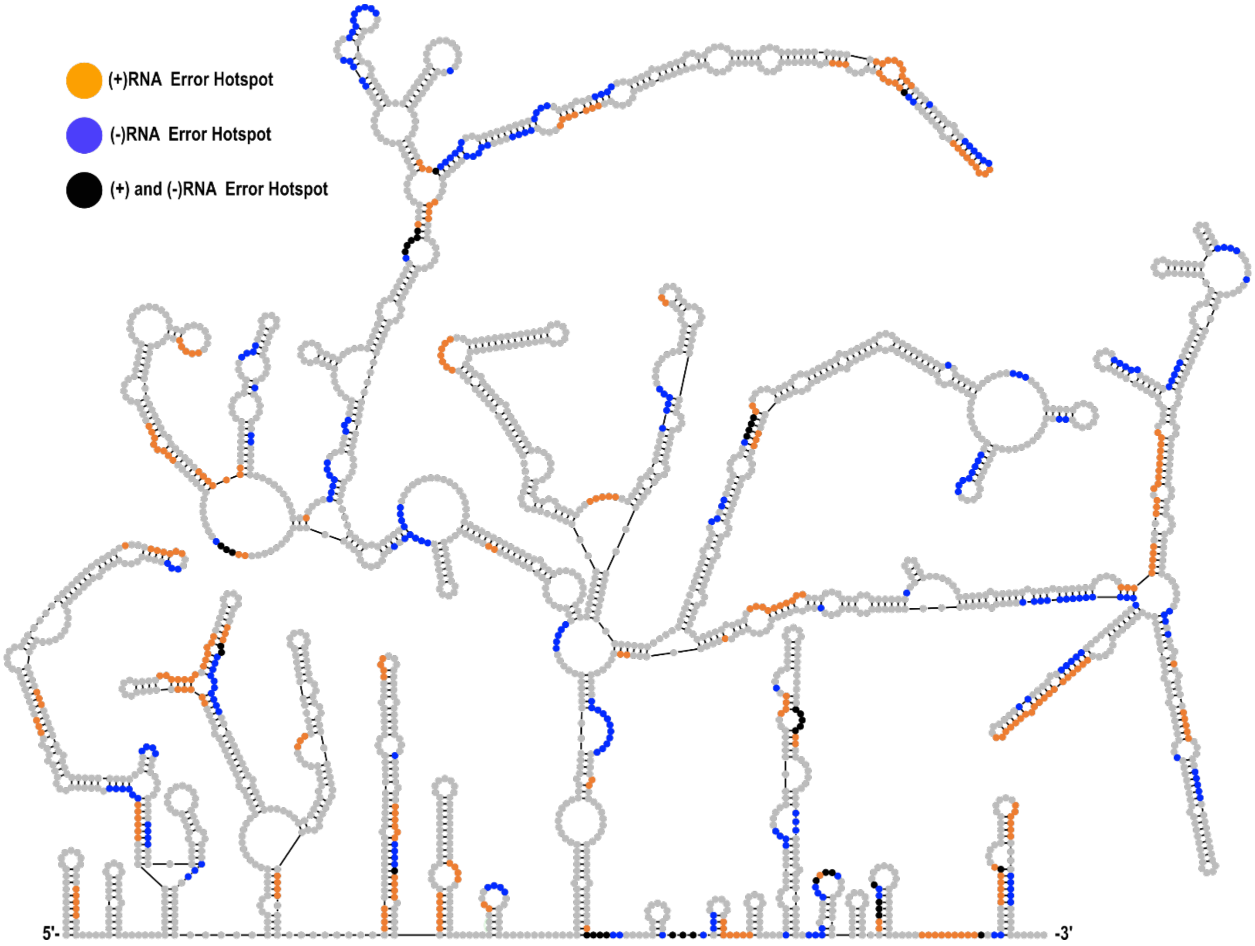


**Supplemental Figure 4. Error hotspots of CY1 (+) and (-)RNA mapped to solved secondary structure.** Reads aligning to either the CY1 (+)RNA (orange) or (-)RNA (blue) were individually aligned to the CY1 reference genome. From these alignments, deletion and mismatch hotspots of were identified by first averaging the deletion or mismatch frequency of a 4-base sliding frame. Hotspots are defined as bases with an error rate greater than 2 standard deviations from the median mismatch or deletion rate calculated from the read subset ([+] or [-]RNA), and were mapped back onto the solved secondary structure of CY1.
